# Using polygenic risk score about complex traits to predict production performance in crossbreeding of yeast

**DOI:** 10.1101/2022.07.07.499257

**Authors:** Yi Dai, Guohui Shi, Mengmeng Chen, Guotao Chen, Qi Wu

**Affiliations:** State Key Laboratory of Mycology, Institute of Microbiology, Chinese Academy of Sciences, Beijing, China; University of the Chinese Academy of Sciences, Beijing, China

**Keywords:** polygenic risk score, hybrid, cross-breeding, complex trait, GWAS

## Abstract

The cultivation of hybrids with favorable complex traits is one of the important goals for animal, plant, and microbial breeding practices. A method that can well predict the production performance of hybrids will be of great significance to the whole research and practice. In our study, polygenic risk scores (PRS) were introduced to estimate the production performance of *Saccharomyces Cerevisiae*. The genetic variation of 971 published isolates and their growth ratios under 35 medium conditions were analyzed by genome-wide association analysis, and the precise *p*-value threshold for each phenotype was calculated. Risk markers for the above 35 phenotypes were obtained. By estimating genotype of F1 hybrids according to that of the parents, the PRS of 613 F1 hybrids was predicted. There was a significant linear correlation between YPD40 and PRS in F1 and their parents (R^2^=0.2582, R^2^=0.2414, respectively), which indicates that PRS can be used to estimate the production performance of individuals and their hybrids. Our method can provide a reference for strains selection and F1 prediction in yeasts cross-breeding, reduce the workload and improve the work efficiency.

## Introduction

Crossbreeding is one of the effective ways to obtain new yeast strains with superior traits. The production of excellent hybrids through cross-breeding has led to a continuous and substantial increase in global agricultural productivity[1–3]. It is usually limited to the complex traits and can be influenced by parental background and imprinting[4]. It facilitates the construction of novel yeast (saccharomyces cerevisiae) strains with preferred characteristics from multiple parental strains by sexual hybridization. Crossbreeding of industrial strains such as baker’s[5], sake[6], and wine yeast strains[7] has been reported. The selection of parents with traits of interests is a prerequisite for obtaining superior hybrid offspring. However, there is no reliable forecasting method for progeny to guide the practice in fungal cross breeding. Many industrial strains have suffered from the low sporulation efficiency and the spore viability[8]. Therefore, it is very meaningful to predict the production performance of the F1 hybrids without extra experiments and time.

Many complex diseases in human beings are caused by both genetic and environmental factors. And most of them are affected by multiple genes[9], so theories of quantitative genetics for complex traits can be used to study such diseases[10, 11]. In the era of omics, high-throughput techniques such as genome-wide genetic association studies (GWAS) have been widely used for comprehensive assessment of genetic susceptibility for various complex traits[12, 13]. In the recent years, the polygenic risk score (PRS) based on GWAS summary results that provide a comprehensive assessment of genetic predisposition for a complex trait, such as height, body weight, cardiovascular disease, and rheumatoid arthritis[14, 15], at the individual level has been widely used in biomedical field. They can effectively identify groups of individuals with substantially increasing risks, so that certain medical treatments or behavioral modifications can be recommended for precaution[14–17]. It weighted the significant risk markers of GWAS to evaluate the genetic liability of individuals to complex traits[18]. That is, PRS is defined as a combination of single nucleotide polymorphisms (SNPs) that associate with the trait of interest[19]. From a statistical point of view, PRS can be considered as a single marker, as similar with an individual biomarker (or biomarker score), has been commonly used for clinically relevant disease prediction[20].

However, PRS is only used to identify the individuals with clinically significant increased risks, or primarily for the early warning of important human traits for adjunctive treatment[15]. In biomedical field, PRS-calculation has now become a popular approach of using GWAS datasets. Even though this approach has not been widely used in non-human organisms, it has been proved as an effective predictor of individuals’ genetic liability to complex traits. In the practice of fungal hybridization, we also pay attention to some complex traits, which have important contributions to production, so whether PRS can be used as an effective indicator of fungal breeding to select high performance yeast strains?

The goal of this study is to identify risk markers about production performance of yeasts, increase the variance explained of phenotypic diversity, and achieve the accurate prediction of phenotypes in F1 hybrids. In this paper, it is proposed to introduce PRS into breeding research, and to screen out high-potential parents based on the parent’s trait performance. In order to predict the phenotype of the next generation of hybrid, we estimated the offspring genotype according to parental genotype, and then estimated its PRS.

## Results

### Identify risk markers for production performance in 35 medium conditions

We performed GWAS analysis on the growth ratios under 35 medium conditions (S1 Fig.). We mainly used two phenotypes of growth ratio at 40 °C in our testing set to verify the relationship between the PRS and the phenotype of the isolates (Fig 2). With p<10e^−8^ as the high-confidence significant threshold, we could find 12 SNPs associated with YPD40 (*R^2^* = 0.33) (Fig 2A and B). When we repeated the analysis using high-resolution PRS and found the most predictive PRS At PT = 0.37885, 2126 SNPs were shown (*R^2^* = 63.97%) (Fig 2C and D). According to LD Score regression results, we also found that LD score regression intercept was close to 1 (0.817±0.0256), indicating that the phenotype was affected by polygenicity.

**Fig 1.**
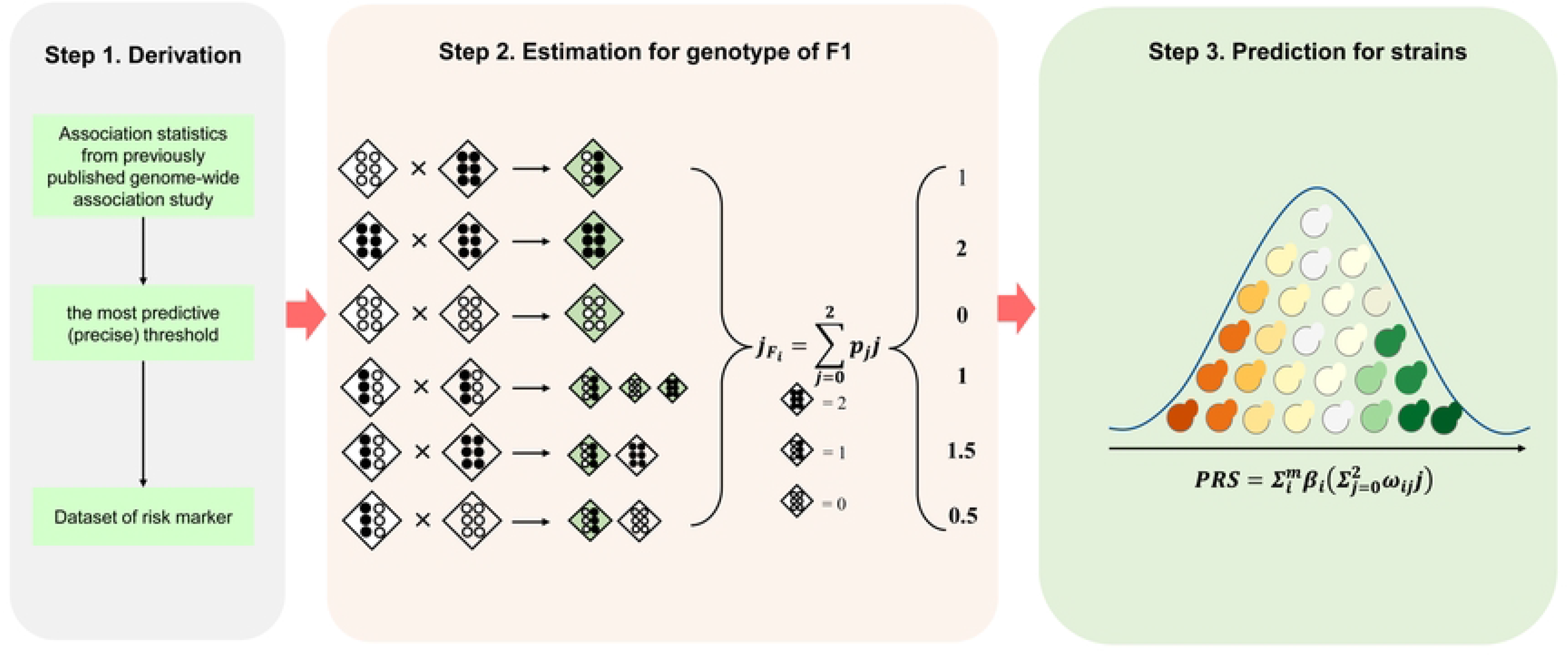
Study design and workflow. We developed a method for predicting the growth ratio of hybrid F1. The method consists of three main steps. First, to identify risk markers associated with the growth ratio, we downloaded the published GWAS summary results. We screened out the risk markers by calculating the precise P value. Then, by estimating the genotypes of F1hybrids according to the parental genotypes, we can calculate their PRS. Finally, the potential of F1 was judged according to the value of PRS.

**Fig 2.**
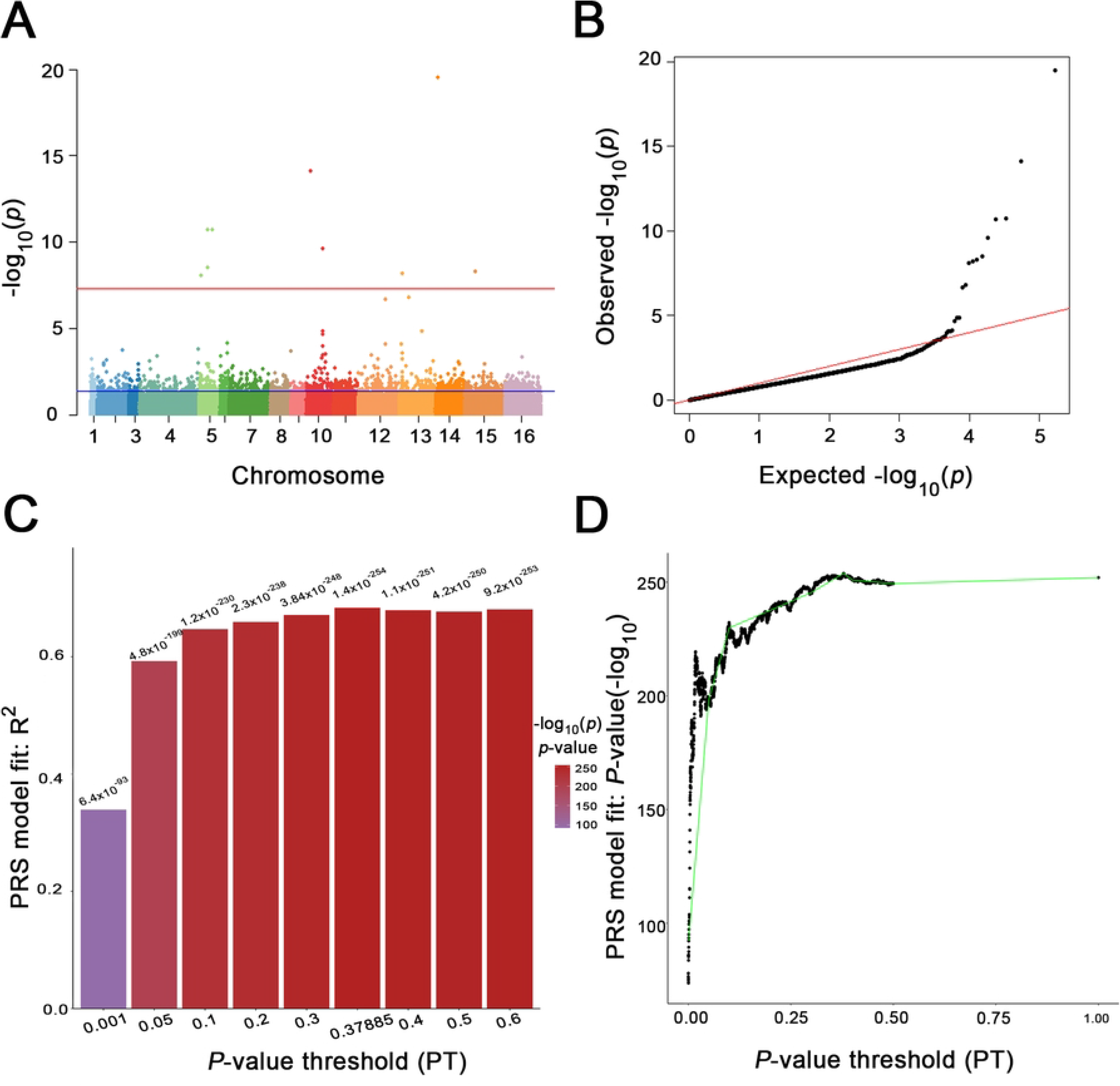
Genome Wide Association Study and Polygenic risk score prediction of YPD40. (A) the Manhattan Plot of GWAS result. The red line indicates that *p*-value is 10e^−8^ and the threshold is statistically significant; the blue line indicates that *p*-value is 0.37885 and the threshold is the precise *p*-value threshold(*p*-value=0.37885). (B) Quantile-Quantile Plot shows good normality. (C) Bar plot from *PRSice-2* showing results at broad *p*-value thresholds for PRS predicting YPD40. A bar for the best-fit PRS from the high-resolution run is also included. (D) High-resolution *PRSice-2* plot for PRS predictingYPD40. The thick line connects points at the broad *p*-value thresholds of (D)

### Relationship between PRSs and phenotypes

There was a significant linear regression between PRS and YPD40 for 52 strains (R^2^=0.2582, *p*-value=1.203e^−4^), indicating that there was a good linear correlation between PRS and phenotype (Fig 3A), which further confirmed that calculating PRS could be used to predict the growth rate of strains at 40°C. At each locus, the combination of parental genotypes and fixed parental loci were analyzed, and they could only produce one kind of gamete. The situations (*AA × AA, AA × aa, aa × aa*) accounted for 99.96% ± 0.04% (Fig 3B). It can be predicted that when both parents are homozygous genotypes, the genotype of the F1 hybrids at this locus can be predicted with relative accuracy. At the same time, we calculated the estimated PRS of F1 by estimating the genotype of F1 hybrids, which also had a good linear relationship and was close to the goodness of fit of parents (Fig 3C).

**Fig. 3.**
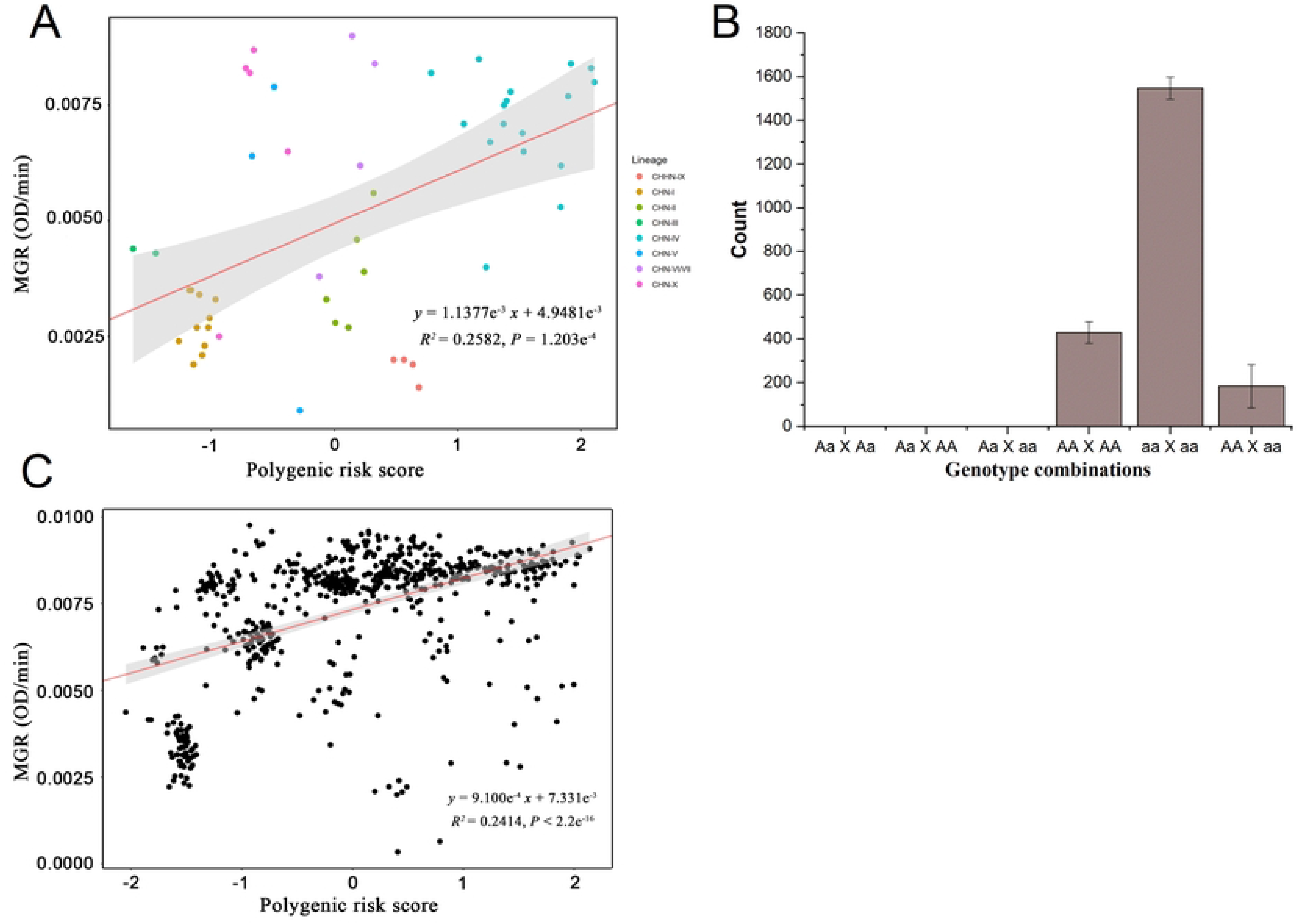
Relation between PRS and YPD40. (A)The linear relationship between PRS and 52 parental strains. (B)Distribution of combinations of genotypes on one locus from two parents of 613 F1 Hybrids. The situations in which both parents were homozygous genotype (*AA* × *AA, aa* × *AA, aa* × *aa*) accounted for 99.96%±0.04%. It can be predicted that when both parents are homozygous genotypes, the genotypes of the F1 hybrids at this locus can be predicted with high accuracy. (C)The linear relationship between PRS and 613 F1 hybrids.

### PRS for heterosis

We found that there was no significant difference in phenotypic distribution among our 52 wild candidate strains compared with other strains, but most of F1 hybrid strains were significantly higher than their parents (Fig 4A), indicating significant heterosis. In terms of PRSs, the distribution of candidate strains and F1 hybrid strains were relatively concentrated, and there was no distribution on the two tails of other strains (Fig 4B). There was a significant difference in PRS between the strains that showed mid-parent heterosis and those that did not (W = 10622, *p*-value = 0.002591), and there was also a significant difference in PRS for the strains that showed depression and other strains (W = 8150, *p*-value = 1.397e^−06^). But there was no significant difference in PRS between the strain with best-parent heterosis and other strains (W = 31429, *p*-value = 0.09518) (Fig 4C). Our results suggest that determine heterosis and depression to a certain extent.

**Fig 4.**
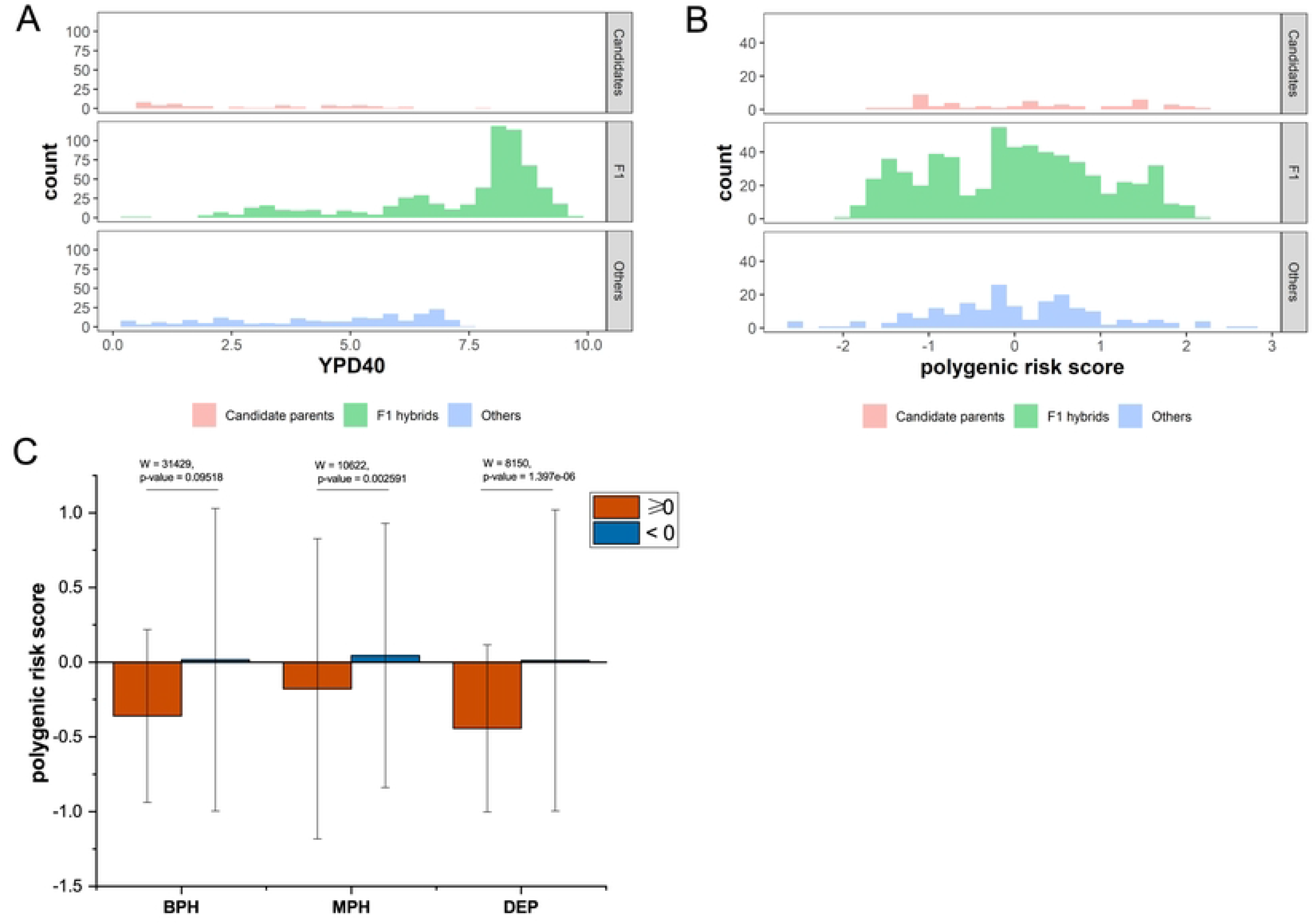
Distribution of YPD40 and PRS of F1 hybrids and 266 isolates for testing set. (A) Distribution of YPD40 of 52 parent strains and 613 F1 hybrids, as well as other strains in the testing set. The phenotype of F1hybrids significantly exceeds that of parents and other strains, suggesting significant heterosis. (B) PRS distribution of 52 parent strains and 613 F1 strains, as well as other strains in the testing set. The parental strain and F1 strain had similar distribution range of PRS. (C) Relationship between heterosis of hybrid strains and PRS. BPH: best-parent heterosis, MPH: mid-parent heterosis, DEP: depression.

## Materials and Methods

We developed a method for predicting the growth ratio of hybrid F1, and validated it with 52 wild-type homozygous *S. cerevisiae* genomes and the growth ratio of their F1 hybrids without any genetic marker through spore-to-spore mating. The method consists of three main steps. First, to identify risk markers associated with the growth ratio, we downloaded the published GWAS summary results about the association between 83794 variant markers and growth ratios under 35 medium conditions. We screened out the risk amrkers by calculating the precise *P* value. Then, by estimating the genotypes of F1hybrids according to the parental genotypes, we can calculate their PRS. Finally, the potential of F1 was judged according to the value of PRS. Fig 1. summarizes the main steps of this process. The Methods section will describe the pipeline in detail.

### Training datasets and testing datasets

To calculate the PRS of one individual, we need to obtain the risk loci associated with each trait. We used a dataset of 1011 *S. cerevisiae* isolates to obtain phenotype association loci. The isolates included in this project were carefully selected from providers and references including 23 kinds of ecological and 312 different geographical origins around the world[21]. Ecology includes the human-related environment as well as the natural environment. The geographic origin is also highly diverse, with a global distribution. These global samples are suitable for us to search for phenotype association loci. Here we use summary statistics from the recent GWAS studies conducted the primarily 971 isolates in the training datasets as well as the phenotypic file about 35 phenotypes (supplement table 1). BED, BIM, and FAM files with all biallelic positions and CNV known for 971 isolates as well phenotypic file were downloaded (http://1002genomes.u-strasbg.fr/files/).

Testing dataset with variants calls of 266 *S. cerevisiae* isolates was a gift from Prof. Fengyan Bai, Institute of Microbiology, Chinese Academy of Sciences. Their work provided the detailed information about the SNPs and the chromosomal CNV of these isolates[22]. They were selected from the different wild lineages which were shown to be homozygous by genome analysis and have the greatest genetic diversity in the wild population of *S. cerevisiae* that has been documented to date[22]. The genome sequence of *S. cerevisiae* S288c reference genome (version R64-1-1) was downloaded from the National Center for Biotechnology Information (NCBI). And the phenotypic data (max growth rate, YPD40) measured for 613 F1 hybrids and their parents without any genetic marker by spore to-spore mating between pairs of the 52 wild *S. cerevisiae* strains gifted from Liang S[23]. (Supplement table 2)

### Identify risk markers

Since the phenotypic data we got were already normalized, and the genomic variants have also been filtered, we can perform the genome-wide association analysis (GWAS) directly. We conducted 971 isolates with MAF>5% as well as CNVs (Peter et al., 2018) to the performed genome-wide association analysis by *GEMMA* 0.98.3 with a linear mixed model and *p*-values from the Wald test to account for 35 phenotypes. And then we computed the variance explained by our significantly associated markers[24] by in-house python script. LD score regression was used to distinguish swelling from true polygenic signals and biases[25]. The method was used after GWAS to quantify the contribution of each factor by examining the relationship between test statistics and linkage disequilibrium (LD). LD score regression intercept can be used to estimate the polygenicity of traits. *PRSice-2*[26] can determine the polygenic risk scores (PRS) under different *p*-value thresholds according to the results of GWAS and provide the best-fit PRS and *p*-value significance threshold. Then, we used it to calculate the PRS of each isolate, and obtained the precise *p*-value threshold (PT), in order to obtain a higher variance explained. Markers whose *p*-value below PT were screened to form a dataset with risk loci associated with the phenotype by in-house python scripts.

### Calculation of the polygenic risk score

PRS were generated by multiplying the genotype dosage of each risk loci by its respective weight, and then summing across all variants in the score using in-house python scripts.

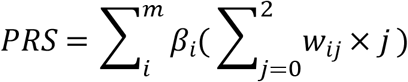

Where *w_ij_* is the probability of observing genotype *j*, where *j*∈ {0,1,2}, for the *i* th SNP; *m* is the number of SNPs; and *β_i_* is the effect size of the *i*th SNP estimated from the relevant GWAS data. The PRS of the candidate parent was calculated based on the distribution of SNPs of the candidate parent. SNPs can be treated as 0/1/2 according to genotype.

### Estimate the genotype of F1 hybrids

For homozygous SNP, there is ideally only one allele that can be passed down to the next generation. But for heterozygous loci, we cannot accurately determine which allele is passed on to the offspring. Assuming that *A* and *a* are two genotypes at one locus, and *A* is the mutant allele, we used 0/1/2 to stably encode *aa, Aa* and *AA* respectively. In this way, we can calculate the mean value of parents at this locus as the estimated genotype of F1. Thus, if both parents are heterozygous at that site, then the offspring are still heterozygous at that site. According to Mendel’s first law of segregation, the probability that the next generation will remain heterozygous in the absence of recombination is the highest (50%). However, for one parent with homozygous genotype and one with heterozygous genotype, our method will produce values of 0.5 and 1.5, which do not exist in the genotype, just for convenience of calculation. Here, we adopted a compromise method, ignoring the possibility of heterozygous site hybridization to produce homozygous, so there is the possibility of underestimation and overestimation.

### Judgment of heterosis of F1 hybrids

To measure the degree of heterosis, F1 was divided into groups according to the above three parameters, and the PRS difference between groups was compared. MPH (Mid-parent Heterosis) > 0 was considered to show heterosis, BPH (Best-parent Heterosis) > 0 was considered to show significant heterosis, and DEP (Depression) > 0 was considered to show decline. The formula is as follows:

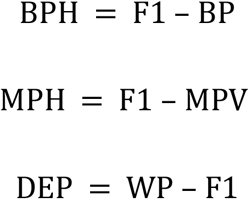

Where F1 is the F1 hybrid phenotype, BP is the maximum of the parental phenotype, MPV is the average of the parental phenotype, WP is the minimum of the parental phenotype.

### Statistical Analysis

the normality was verified by Kolmogorov-Smirnov test. we perform linear fitting on the PRS and phenotype data of 52 candidate parents and 613 F1 hybrids, and output 95% confidence interval. All tests were two-tailed with alpha threshold of 0.05. Statistical analyses were conducted in R v3.6.0.

### Code availability

The scripts for calculating PRS for parents, gametes, and any F1 hybrids were deposited in GitHub https://github.com/DYqwert/pyprs/

## Discussion

Achieving trait prediction of F1 hybrid is of great significance to cross-breeding, whether in terms of industrial, experimental costs, or genetic research. Our method has enabled the efficient prediction of traits in the progeny of hybrids, and confirmed the correlations with the experimental results. This method can accurately predict the phenotype of hybrid generation only by parental phenotype and genome and hybridization experiments are not conducted. Moreover, this method is not only applicable to the production traits mentioned in this paper, but also to other important complex traits (S2 Fig.). It is hoped that this method can be helpful for the crossbreeding of fungi.

PRS are similar to narrow heritability in that they represent the aggregate and additive effects of segregating loci of small effect to some degree[27, 28]. Considered as a quantification of the additive effect of an individual’s risk genes, PRS can be used to estimate the additive genetic variance as a hybrid offspring. In the process of gene transmission, the additive genetic effects are relatively stable. Therefore, it is feasible and meaningful to estimate the productivity of the hybrids by evaluating the potential of their parents based on the multiple variants related to productivity. While prediction of phenotype from an individual’s genetic profile is compromised by this polygenicity, the application of PRS has shown that prediction is sufficiently accurate for several applications[17, 18]. Our method also unlike genomic selection, which refer to the use of dense markers covering the whole genome to estimate the breeding value of selection candidates for a quantitative trait[29, 30]. With the risk loci selected by the high-resolution PRS, the method can reduce the false negatives on GWAS results and the false positives on genomic selection results, furthermore significantly increase the PVE, and has a significant improvement on screening efficiency without progeny testing. Unlike the previous application of precision medicine and preventive medicine[15], we can use PRS in our method in breeding research to extend its application on non-human organisms’ prediction.

For homozygous SNP, ideally only one allele can be inherited to the next generation. But for heterozygous loci, we cannot accurately determine which allele is passed on to the offspring. Our method may underestimate or overestimate the effect of homozygous loci. Simply put, under the assumption that there are only two alleles, the offspring can only have three genotypes. Considering the amount of calculation, we only need to estimate the minimum and maximum PRS of different F1 genotypes, and the obtained interval can be used as a breeding reference. Theoretically, the variance of PRS mainly comes from the genotype difference produced by uncertain gametes, so with the increase of uncertain genotypes, the variance of estimated PRS will increase. In addition, our method yet cannot predict the heterosis of hybrids. However, our results show that PRS may be used to judge whether heterosis exists, although it cannot be used to judge the degree of heterosis. Heterosis is an important reason for cross-breeding research, so the prediction of F1 heterosis is also crucial. However, the complementation of the allelic variation and the variation in gene content and gene expression patterns are likely to be an important contributor to heterosis[4]. There are some evidences that have shown the possible roles of nonadditive genes in manifestation of heterosis or outbreeding the depression in *S. cerevisiae* [23]. Although the additive effect and the epistatic effect have certain mutual influence[31], it is difficult to estimate the phenotype difference between hybrids and their parents caused by non-additive effects only by the PRS of hybrids. Generally, the additive effect is relatively stable. We have been able to predict the narrow-sense heritability efficiently, but the prediction for broad-sense heritability alone is unreliable with PRS. So, how to accurately predict the broad-sense heritability will be the focus of our further research.

## Acknowledgements

We thank Prof. Fengyan Bai and Dr. Liang Song for their gifts of yeast strains and phenotypic data. This research is financially supported by the National Natural Science Foundation of China (Grant No.32170015), the Strategic Priority Research Program of Chinese Academy of Sciences (grant No. XDB31000000 and XDA28030401), the National Natural Science Foundation of China (Grant No. 91746119), and the Senior User Project of RV KEXUE (grant no. KEXUE2019GZ05).

## Supporting information

**S1 Fig. Genome Wide Association Study of 1011 isolates under 35 conditions.**

The red line indicates that p-value is 10e^−8^ and the threshold is statistically significant; the blue line indicates that p-value is 0.37885 and the threshold is the precise p-value threshold.

**S2 Fig. The distribution of polygenic risk score of 266 strains in testing set.**

**S1 Table. Growth conditions tested for phenotyping in training set.**

